# ALS-associated mutations reveal a conserved C-terminal self-assembly module in the TDP-43 low-complexity domain

**DOI:** 10.64898/2026.10.09.756561

**Authors:** Nobuo Maita, Sun-Yong Kim, Yuko Kajino, Masato Kato

## Abstract

TAR DNA-binding protein 43 (TDP-43) is an RNA-binding protein whose low-complexity domain (LCD) mediates self-assembly, liquid-liquid phase separation, and pathological aggregation associated with amyotrophic lateral sclerosis (ALS) and frontotemporal dementia (FTD). Previous studies have established the central ultraconserved region (UCR) of the LCD as a major determinant of TDP-43 self-association. However, the contribution of the evolutionarily conserved C-terminal region remains poorly understood. To identify LCD residues required for self-assembly, we systematically analyzed 47 ALS-associated substitutions distributed throughout the TDP-43 LCD using hydrogel-binding and thioflavin T (ThT) assays. Several ALS-associated substitutions within the C-terminal conserved region (CCR) markedly reduced hydrogel incorporation and impaired fibrillization. Similar assembly defects were observed for methionine-to-serine substitutions at two conserved CCR residues, M405 and M414. Furthermore, CCR-containing C-terminal fragments encompassing residues 360–414 or 379–414 formed ThT-positive fibrillar assemblies *in vitro*. The assembly kinetics exhibited a characteristic lag phase followed by rapid fluorescence growth, consistent with the formation of cross-β-rich assemblies. These findings identify the CCR as an additional conserved self-assembly module within the TDP-43 LCD and expand the prevailing UCR-centered model of TDP-43 assembly. The observation that multiple ALS-associated substitutions reduce rather than enhance self-association further raises the possibility that disruption of normal TDP-43 assembly contributes to disease pathogenesis.

## Introduction

TAR DNA-binding protein 43 (TDP-43) is a highly conserved, ubiquitously expressed nuclear RNA-binding protein that plays a central role in various aspects of RNA metabolism, including transcriptional regulation, alternative splicing, mRNA transport, and stability (Guo et al. 2017). TDP-43 was originally identified as a protein that binds the trans-activation response element of HIV-1. It subsequently emerged as a central protein in neurodegeneration following its identification as the major component of ubiquitin-positive inclusions in amyotrophic lateral sclerosis (ALS) and frontotemporal lobar degeneration (FTLD) (Neumann et al. 2006).

Human TDP-43 has 414 amino acids composed of several functional domains (Figure 1A). The N-terminal domain (NTD) mediates self-association and oligomerization, which are important for normal RNA processing. A nuclear localization signal (NLS) near the N-terminus directs TDP-43 to the nucleus, where it predominantly functions. TDP-43 contains two conserved RNA recognition motifs (RRM1 and RRM2) that bind UG-rich RNA and DNA sequences. These domains enable TDP-43 to regulate diverse aspects of RNA metabolism, with RRM1 providing strong RNA affinity and RRM2 contributing to specificity (Guo et al. 2017).

**Figure 1.**
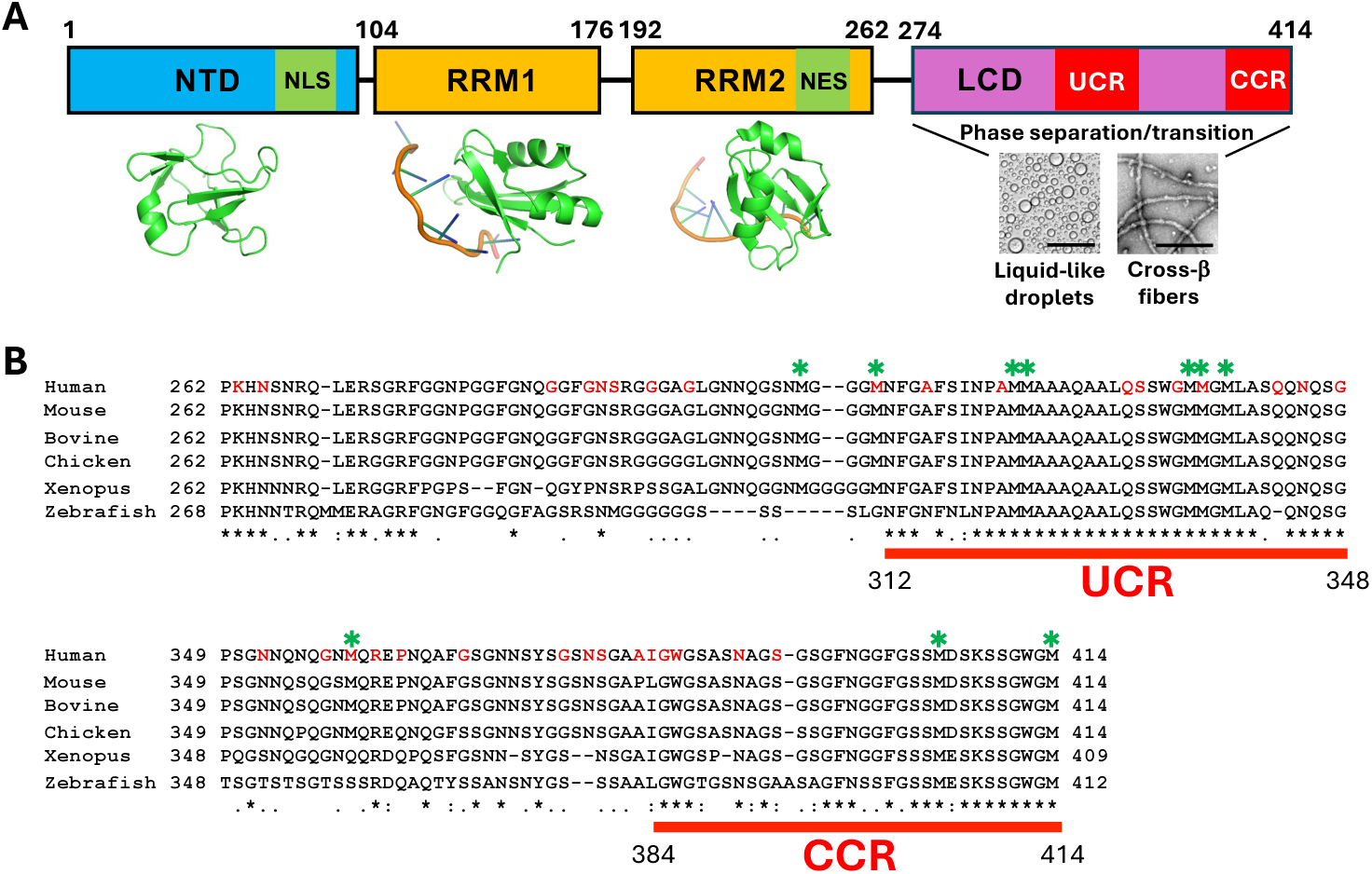
Domain organization and conserved regions of TDP-43. **(A)** Domain organization of human TDP-43. TDP-43 consists of an N-terminal domain (NTD), nuclear localization signal (NLS), nuclear export signal (NES), two RNA-recognition motifs (RRM1 and RRM2), and a C-terminal low-complexity domain (LCD). Solution NMR or crystal structures of each domain except the LCD are shown below the diagram; NTD: PDB 5X4F (Jiang et al. 2017), RRM1: PDB 4IUF (Kuo et al. 2014), RRM2: PDB 3D2W (Kuo et al. 2009). The LCD mediates liquid-liquid phase separation and cross-β fiber formation. The residue numbers are based on the human TDP-43 sequence. **(B)** Sequence conservation of the two evolutionarily conserved regions within the LCD. Multiple sequence alignment of vertebrate TDP-43 proteins identifies the central ultraconserved region (UCR; residues 312-348) and the C-terminal conserved region (CCR; residues 384-414). Green asterisks shown above the sequence indicate conserved methionine residues. ALS-associated mutations analyzed in this study are colored in red in the human sequence. The residue numbers for UCR and CCR are based on the human TDP-43 sequence.

The C-terminal low-complexity domain (LCD) has a strong propensity for self-assembly and mediates reversible liquid–liquid phase separation (LLPS), enabling the formation of dynamic ribonucleoprotein granules under physiological conditions (Kato et al. 2012; Molliex et al. 2015). Under certain conditions, the LCD can also assemble into amyloid-like fibrils containing cross-β structure (Jiang et al. 2013; Babinchak et al. 2020). Cellular stress, disease-associated mutations, and other perturbations have been proposed to shift TDP-43 assemblies toward more ordered and less dynamic states. Cross-β interactions may contribute both to reversible LCD assembly and to the formation of pathological TDP-43 aggregates observed in neurodegenerative diseases, including ALS and FTLD (Kato et al. 2012; Molliex et al. 2015), although the precise relationship between these assembly states remains incompletely understood.

Within the LCD of TDP-43, there are two sequence regions that are evolutionarily conserved: the central ultra-conserved region (UCR, residues 312–348) and the C-terminal conserved region (CCR, residues 384–414) (Figure 1B). The UCR has been shown to be a major determinant of LCD self-association. This region transiently forms an α-helix in solution when it is in a monomeric state (Jiang et al. 2013). This α-helix can intermolecularly interact with the same α-helix of other TDP-43 LCD molecules, which is proposed to be a driving force of phase separation (Conicella et al. 2016). Depending on the oxidation state of conserved methionines in this region, as well as the pH and salt concentration of the solution, the UCR can transform into a β-strand conformation concomitant with interactions with the corresponding β-strand of other TDP-43 LCD molecules, which has also been proposed to play a role in phase separation and cross-β fiber formation (Conicella et al. 2016; Zhou et al. 2022; Gu et al. 2023). This oxidation-dependent β-to-α conformational switch is proposed to trigger the dissociation of TDP-43 from cross-β-rich RNA granules and its subsequent association with the mitochondrial membrane (Gu et al. 2023).

The CCR (residues 384–414) is located at the extreme end of the TDP-43 LCD (Figure 1). This region contains several ALS-associated mutations and the pathological phosphorylation sites S403/S404 and S409/S410. Hyperphosphorylation of these residues is consistently observed in TDP-43-positive inclusions from patients with ALS, frontotemporal dementia (FTD), and other TDP-43 proteinopathies, making them hallmark pathological signatures of disease-associated TDP-43 (Hasegawa et al. 2008; Gruijs da Silva et al. 2022). The strong evolutionary conservation of this region further supports its importance in TDP-43 function and proteostasis. Although the CCR contains multiple disease-associated mutations and major pathological phosphorylation sites, its contribution to TDP-43 self-association and higher-order assembly remains poorly defined.

In this study, we examined 47 ALS-associated substitutions distributed throughout the TDP-43 LCD, together with individual methionine-to-serine substitutions at its ten conserved methionine residues. Hydrogel-binding and Thioflavin T (ThT) assays identified multiple substitutions within the CCR that impaired LCD self-assembly. Furthermore, short CCR-containing C-terminal fragments formed ThT-positive fibrillar assemblies *in vitro*. These findings identify the conserved C-terminal region as an additional self-assembly module within the TDP-43 LCD.

## Results

### Design and purification of fluorescently tagged TDP-43 LCD constructs

Initial constructs in which mCherry or GFP was fused directly to the N terminus of the TDP-43 LCD were robustly expressed in *Escherichia coli* (Kato et al. 2012; Kato et al. 2017), but the resulting cultures and purified proteins occasionally exhibited little or no visible fluorescence. A similar loss of fluorescence has been observed for fluorescent proteins fused to strongly self-associating LCDs, presumably because rapid or extensive LCD assembly interferes with fluorophore maturation or stability (Gu et al. 2025). To minimize this effect, mCherry or GFP was repositioned at the C terminus of the TDP-43 LCD and separated from the LCD by a flexible (GGSG)_5_ linker, which consists of five tandem repeats of the GGSG tetrapeptide (ln5). The resulting TDP-43 LCD-ln5-mCherry/GFP constructs reproducibly retained fluorescence following bacterial expression and purification. Using this configuration, we generated and purified GFP-tagged LCD constructs containing ALS-associated substitutions or individual methionine-to-serine substitutions (Supplementary Figures 1 and 2).

**Figure 2.**
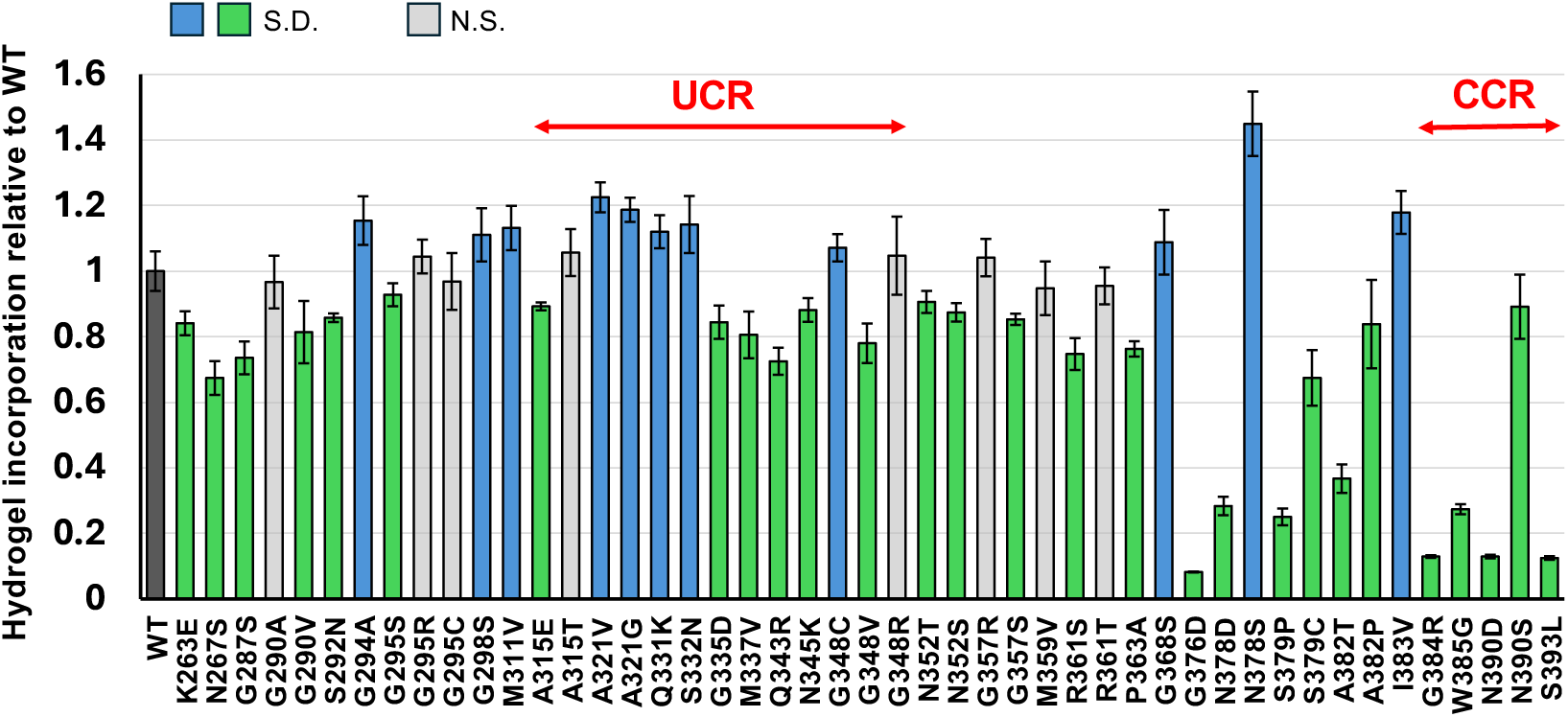
Hydrogel-binding analysis of ALS-associated mutants. Hydrogel-binding activity of GFP-tagged ALS-associated TDP-43 LCD mutants was quantified using preformed mCherry-tagged TDP-43 LCD hydrogels. Fluorescence intensity within hydrogel droplets was normalized to that of wild-type TDP-43 LCD. Data represent the mean ± standard deviation from three independent experiments. Mutants exhibiting significantly higher fluorescence intensities than the wild type are shown in blue, whereas mutants exhibiting significantly lower fluorescence intensities are shown in green. Statistical significance was determined by one-way ANOVA followed by Dunnett’s multiple-comparisons test using the wild type as the control. S.D., significantly different from wild type (P < 0.05); N.S., not significant.

### Hydrogel-binding assays of ALS-associated TDP-43 LCD mutants

In the hydrogel-binding assay, preformed hydrogel droplets generated from wild-type TDP-43 LCD-ln5-mCherry were incubated with wild-type or mutant TDP-43 LCD-ln5-GFP. GFP signal accumulation within the mCherry-labeled hydrogel droplets was subsequently examined by confocal fluorescence microscopy. Signal accumulation primarily reflects the incorporation of TDP-43 LCD-ln5-GFP monomers (test protein) onto the ends of the existing cross-β fibers (seed fibers) of TDP-43 LCD-ln5-mCherry in the hydrogel (Kato et al. 2017). Therefore, this assay reflects the ability of a test protein to elongate the seed fibers. Consequently, the GFP signal intensity within the hydrogel reflects the capacity of the test protein to form cross-β fibers.

The GFP signals of ALS-associated TDP-43 LCD mutants incorporated into hydrogel droplets varied considerably among mutants (Supplementary Figure 3). We quantified the mean GFP intensity of each mutant from three independent experiments and compared it with that of the wild-type LCD (Figure 2). Although many mutants showed statistically significant differences from the wild type, the overall changes were relatively modest, ranging approximately from 0.8-to 1.2-fold of the wild-type value. Several mutations located in the UCR, including A321V, A321G, Q331K, S332N, and G348C, exhibited significantly higher GFP intensities than the wild type (blue bars), suggesting enhanced incorporation into cross-β fibers of TDP-43 LCD. In contrast, a substantial number of mutants displayed significantly reduced GFP intensities relative to the wild type across the LCD sequence (green bars). Notably, the most pronounced reductions were observed in the C-terminal region, including the CCR (Figure 2). All five ALS-associated mutations in the CCR (G384R, W385G, N390D, N390S, and S393L) significantly decreased GFP incorporation into the hydrogels. These results suggest that CCR mutations impair the ability of TDP-43 LCD to assemble into cross-β fibers, representing loss-of-assembly mutations. In contrast, previous studies have primarily characterized ALS-associated mutations that enhance fibril formation by TDP-43 LCD, exemplified by several mutations in the UCR (Lim et al. 2016; Zhou et al. 2022). Therefore, to better understand disease mechanisms associated with reduced assembly, we focused subsequent analyses on the CCR mutations.

### Thioflavin T assays for the ALS mutants of TDP-43 LCD

To validate the results of the hydrogel-binding assay using an independent approach, we performed Thioflavin T (ThT) assays on ALS-associated mutations located in the C-terminal region of the LCD, including the CCR (Figure 3 and Supplementary Figure 4). Fibrillization kinetics were evaluated by measuring the lag time, defined as the interval between the start of the assay and the onset of the increase in ThT fluorescence (see Materials and Methods). Compared with the wild-type protein, many ALS mutants exhibited similar or prolonged lag times. Notably, four of the five CCR mutants (W385G, N390D, N390S, and S393L) showed significantly extended lag times or failed to exhibit a detectable increase in ThT fluorescence during the 24-h observation period, indicating markedly delayed or completely abolished fibrillization. Consistent with the hydrogel-binding assay, these four mutants also exhibited significantly reduced incorporation into hydrogel droplets (Figure 2). Together, these results demonstrate that these CCR mutations substantially impair the ability of TDP-43 LCD to assemble into amyloid-like fibers.

**Figure 3.**
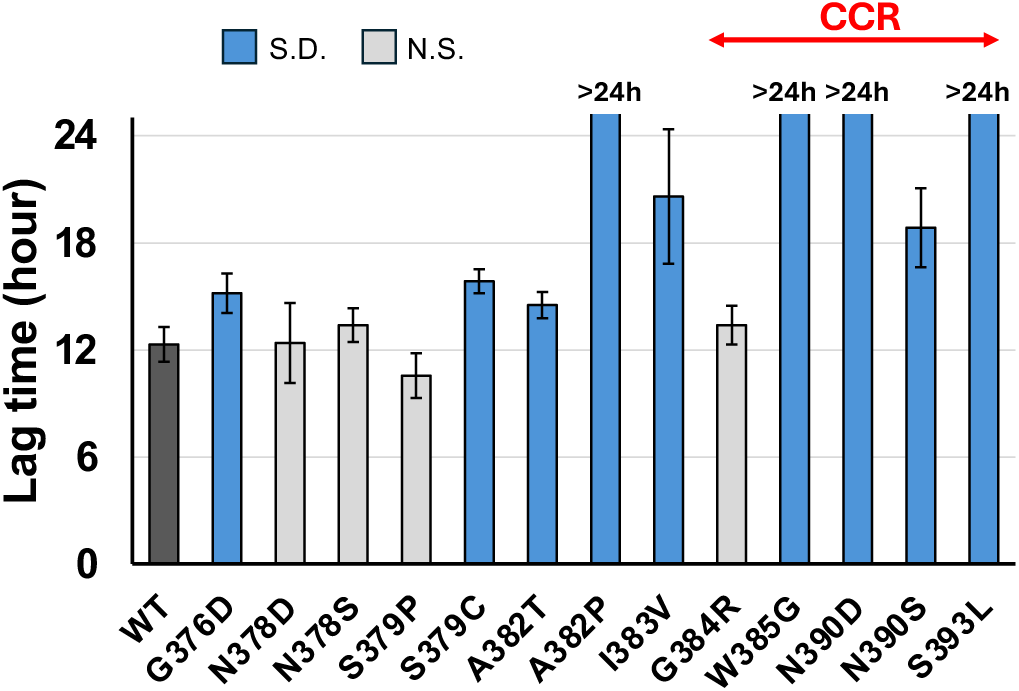
ALS-associated mutations within the CCR alter the fibrillization kinetics of the TDP-43 LCD. Lag times of wild-type and ALS-associated mutant TDP-43 LCD proteins were determined from the corresponding ThT fluorescence traces (Supplementary Figure 4) as described in Materials and Methods. Data are presented as the mean ± standard deviation from six replicate wells. Mutants exhibiting significantly longer lag times than the wild type are shown in blue. Samples that did not reach the lag-time criterion during the 24-h observation period are indicated as “>24 h”. Statistical significance was determined by Kaplan-Meier survival analysis to account for right-censored data (>24 h), followed by the log-rank test with Bonferroni correction. S.D., significantly different from wild type (P < 0.05); N.S., not significant.

### Hydrogel-binding and ThT assays of TDP-43 LCD methionine-to-serine mutants

TDP-43 LCD contains 10 highly conserved methionines (Figure 1B). Oxidation of these methionines was reported to increase the solubility of TDP-43 LCD, resulting in the dissolution of liquid-like droplets or the disassembly of cross-β fibers (Lin et al. 2020). Because the reported solubility of methionine sulfoxide is comparable to that of serine (Wolschner et al. 2009), methionine-to-serine substitutions were used as a simple approximation of the increase in side-chain polarity associated with methionine oxidation. Accordingly, we constructed 10 mutants in which each methionine was individually replaced with serine (Supplementary Figure 2). We evaluated these mutants using hydrogel-binding and ThT assays.

In the hydrogel-binding assays, many mutants exhibited binding similar to or significantly greater than that of the wild type (Figure 4 and Supplementary Figure 5). In contrast, two mutants (M405S and M414S) in the CCR exhibited significantly reduced binding. In the ThT assays, these two mutants also exhibited longer lag times than the wild type (Figure 5 and Supplementary Figure 6). Together, these findings identify M405 and M414 in the CCR as the only methionine residues whose substitution consistently impaired TDP-43 LCD self-assembly in both assays.

**Figure 4.**
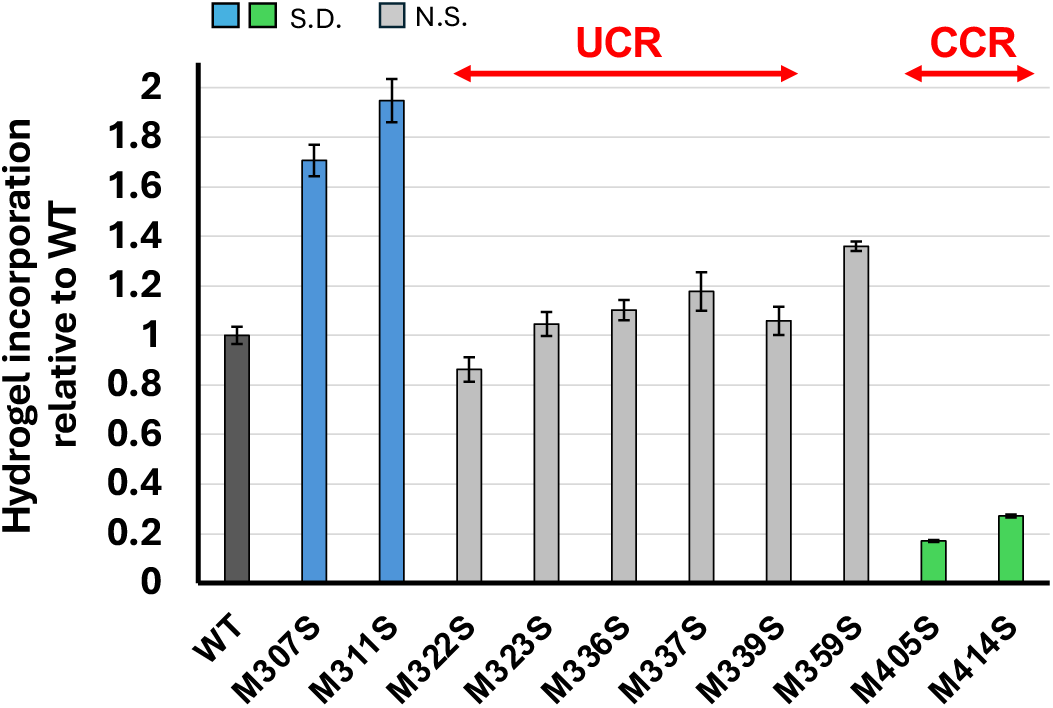
Hydrogel-binding assays of methionine-to-serine substitutions of TDP-43 LCD. Hydrogel-binding activity of methionine-to-serine (M-to-S) mutants of TDP-43 LCD was quantified using preformed mCherry-tagged TDP-43 LCD hydrogels. Fluorescence intensity within hydrogel droplets was normalized to that of wild-type TDP-43 LCD. Data represent the mean ± standard deviation from three independent experiments. Mutants exhibiting significantly higher fluorescence intensities than the wild type are shown in blue, whereas mutants exhibiting significantly lower fluorescence intensities are shown in green. Statistical significance was determined by one-way ANOVA followed by Dunnett’s multiple-comparisons test using the wild type as the control. S.D., significantly different from wild type (P < 0.05); N.S., not significant.

**Figure 5.**
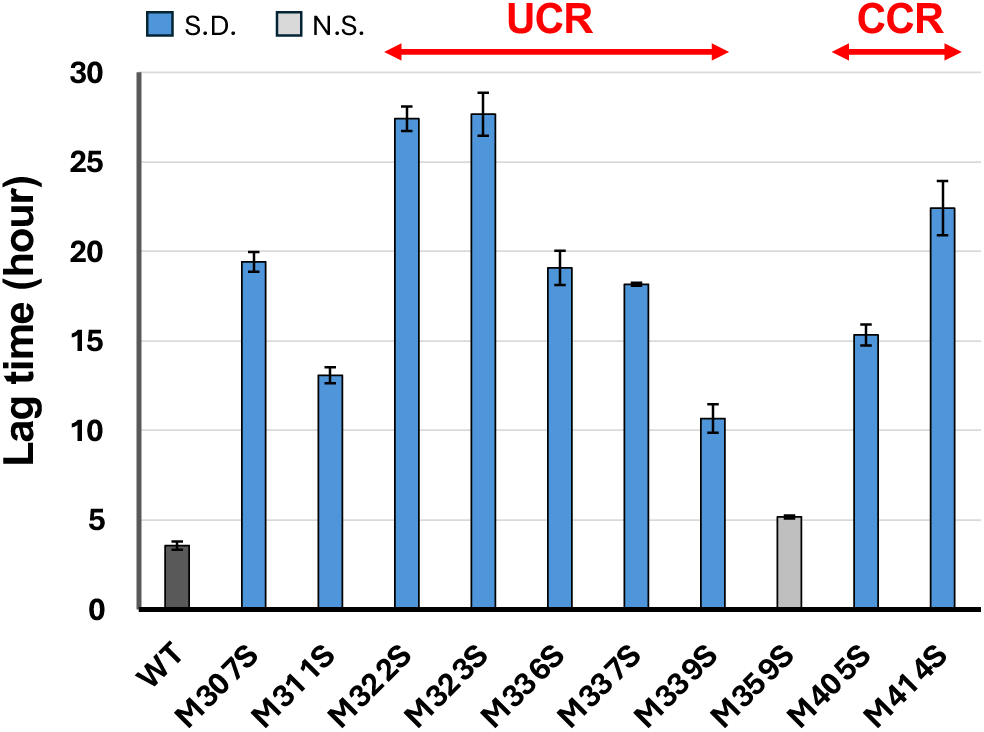
Effects of methionine-to-serine substitutions on the fibrillization kinetics of the TDP-43 LCD. Lag times of wild-type and methionine-to-serine mutant TDP-43 LCD proteins were determined from the corresponding ThT fluorescence traces (Supplementary Figure 6) as described in Materials and Methods. Data are presented as the mean ± standard deviation from six replicate wells. Mutants exhibiting significantly longer lag times than the wild type are shown in blue. Statistical significance was determined by one-way ANOVA followed by Dunnett’s multiple-comparisons test using the wild type as the control. S.D., significantly different from wild type (P < 0.05); N.S., not significant.

### A CCR-containing C-terminal fragment is sufficient to form fibrillar assemblies

To examine the role of the CCR in TDP-43 LCD self-association, we purified C-terminal truncation constructs encompassing residues 360-414 or 379-414 and induced assembly *in vitro*. Both fragments exhibited robust increases in ThT fluorescence following a lag phase, producing sigmoidal assembly kinetics characteristic of fibril growth (Figure 6B). Negative-stain electron microscopy revealed abundant unbranched fibrillar structures in both samples (Figure 6C). The agreement between the ThT and electron microscopy analyses indicates that these fragments form ThT-positive fibrillar assemblies consistent with cross-β-rich structures. Together, these findings demonstrate that short CCR-containing C-terminal segments possess an intrinsic capacity for self-assembly even in the absence of upstream LCD regions, including the UCR.

**Figure 6.**
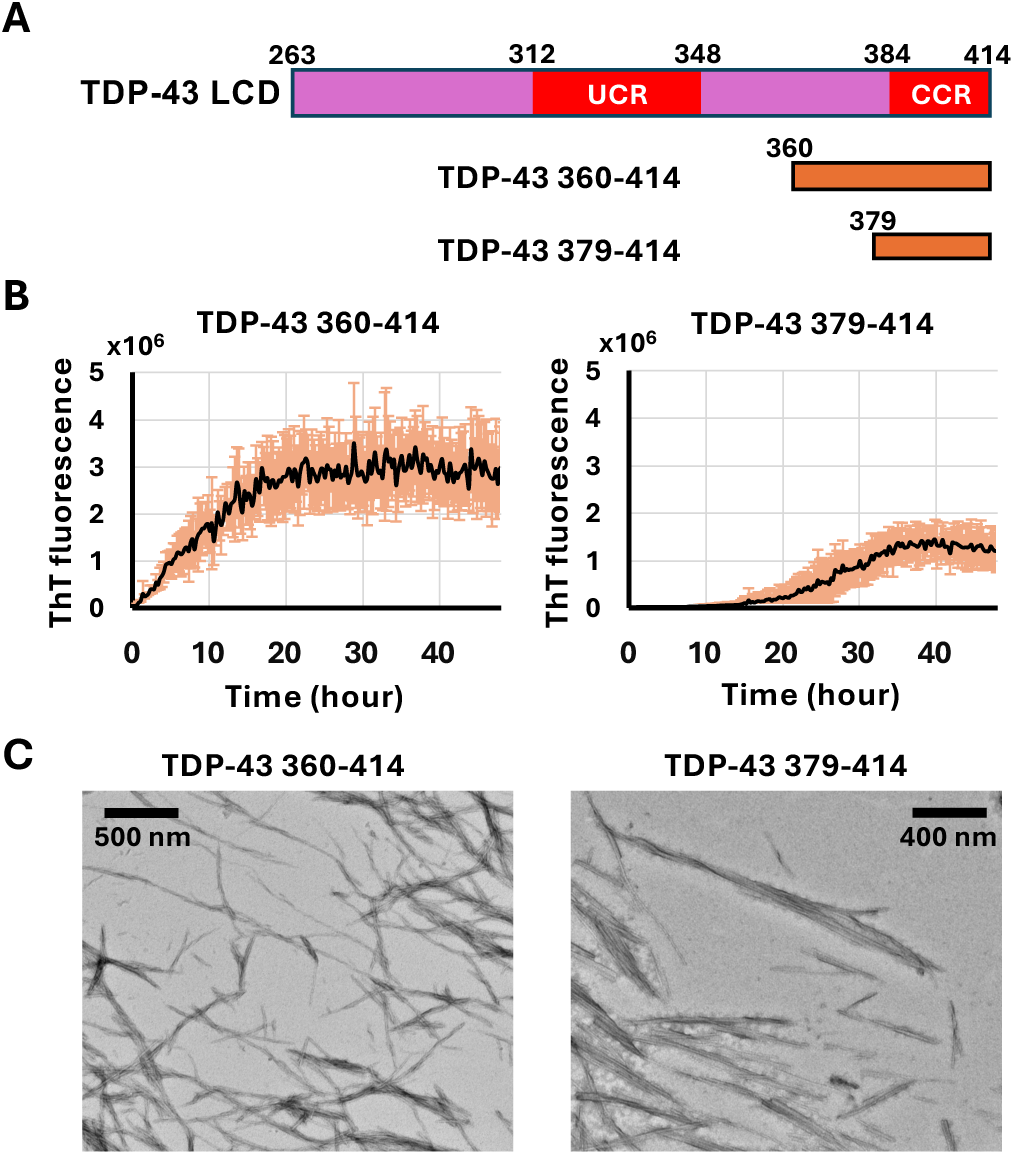
CCR-containing C-terminal fragments undergo ThT-positive fibrillar assembly *in vitro*. **(A)** Schematic representation of the TDP-43 C-terminal fragments encompassing residues 360-414 and 379-414 used for fibrillization experiments. The location of the CCR is indicated in red. **(B)** Representative ThT fluorescence traces of the 360-414 and 379-414 fragments under fibrillization conditions. Both fragments exhibited a lag phase followed by a rapid increase in fluorescence intensity, consistent with fibril growth. Each trace represents the mean fluorescence intensity obtained from six replicate wells. The orange bars indicate standard deviation. **(C)** Negative-stain electron micrographs of assemblies formed by the two C-terminal fragments after incubation under fibrillization conditions. Numerous unbranched fibrillar structures were observed. Scale bars, 500 nm for the left image and 400 nm for the right image.

## Discussion

### The CCR defines an additional C-terminal self-association module

Previous studies have identified the central ultraconserved region (UCR) of the TDP-43 LCD, particularly residues 312-348, as a major determinant of self-association for liquid–liquid phase separation and cross-β interaction (Conicella et al. 2016; Zhou et al. 2022). ALS-associated mutations within this region alter intermolecular contacts and modulate phase separation behavior, supporting the view that the UCR serves as a key assembly module of TDP-43.

Our findings reveal that the C-terminal conserved region (CCR, residues 384-414) also plays a critical role in TDP-43 self-assembly. Several ALS-associated mutations (G384R, W385G, N390D, and S393L) and two methionine-to-serine substitutions (M405S and M414S) located within the CCR substantially reduced hydrogel incorporation and fibril formation (Figures 2-5). Furthermore, CCR-containing fragments (360-414 and 379-414) underwent ThT-positive fibrillization and formed unbranched fibrils *in vitro* (Figure 6), demonstrating that the CCR possesses an intrinsic capacity for self-association independent of the UCR. The more rapid assembly of the 360-414 fragment further suggests that residues immediately upstream of the CCR may facilitate this process, consistent with previous structural observations of aged TDP-43 assemblies (Fonda et al. 2021).

### A revised model: coordinated contributions of the UCR and CCR

Our results suggest that the current model of TDP-43 assembly, in which the UCR acts as the principal assembly determinant, may be incomplete. The identification of multiple assembly-impairing mutations within the CCR, together with the intrinsic ThT-positive fibrillization capacity of the 360-414 and 379-414 CCR-containing fragments, indicates that at least two conserved regions contribute to TDP-43 LCD assembly. The presence of two assembly regions may represent a broader organizational principle of RNA-binding protein LCDs. The FUS LCD similarly contains distinct N- and C-terminal cross-β cores that have been proposed to function in a mutually exclusive manner, thereby balancing alternative modes of self-association (Kato et al. 2021). Whether the UCR and CCR of TDP-43 cooperate, compete, or become preferentially engaged under different conditions remains to be determined.

### Specific importance of M405 and M414

Methionine oxidation has been shown to disrupt TDP-43 LCD self-association and phase separation (Lin et al. 2020; Carrasco et al. 2023). A recent NMR and molecular simulation study further demonstrated that oxidation impairs TDP-43 CTD phase separation through region-dependent mechanisms: oxidation within the conserved helical region reduces helicity and disrupts helix-mediated association, whereas oxidation within the flanking disordered regions weakens transient long-range contacts (Ozguney et al. 2026).

Our findings extend these observations by identifying two CCR methionines, M405 and M414, as particularly important for self-assembly (Figures 4 and 5). Although methionine-to-serine substitutions do not fully reproduce methionine oxidation, the selective effects of M405S and M414S support a direct contribution of the CCR to TDP-43 assembly.

### ALS mutations do not always promote aggregation

Pathogenic TDP-43 mutations have frequently been interpreted within a gain-of-function framework in which disease-associated substitutions enhance aggregation propensity and promote the formation of pathological inclusions. However, accumulating evidence indicates that some ALS-associated mutations instead impair physiologically relevant intermolecular interactions and phase separation (Conicella et al. 2016; Guo et al. 2017; Conicella et al. 2020; Gruijs da Silva et al. 2022).

Consistent with this view, most of the ALS mutations in the CCR examined here markedly reduced hydrogel incorporation and delayed or abolished ThT-positive fibrillization (Figures 2 and 3). These findings demonstrate that pathogenic mutations can decrease rather than increase the self-assembly potential of the TDP-43 LCD.

Because TDP-43 self-association contributes to normal biological functions, including phase separation and RNA processing, disruption of physiological assembly may itself represent a disease-relevant mechanism (Mann et al. 2019; Gruijs da Silva et al. 2022). Therefore, TDP-43-associated neurodegeneration should not be viewed solely through the lens of aggregation-promoting mutations but also through mutations that compromise normal assembly behavior.

### Relationship between CCR and pathological phosphorylation

The CCR contains the major disease-associated phosphorylation sites S403/S404 and S409/S410 (Hasegawa et al. 2008; Kametani et al. 2016). Because several ALS-associated mutations and methionine substitutions within the same region impaired self-association (Figures 2-5), the CCR may serve as a regulatory hub through which post-translational modifications modulate TDP-43 assembly. Recent studies indicate that phosphorylation alters condensation and fibril properties of TDP-43 LCD (Haider et al. 2024; Fonda et al. 2026), supporting the possibility that phosphorylation influences CCR-mediated interactions. Future studies will be required to determine how phosphorylation affects CCR-dependent assembly and whether these mechanisms contribute to disease-associated TDP-43 states.

## Materials and Methods

### Cloning, expression and purification of proteins

Cloning of the expression plasmids encoding N-terminally His-tagged and His-mCherry/GFP-tagged TDP-43 LCD (residues 263-414) has been described previously (Kato et al. 2012). Expression plasmids encoding TDP-43 LCD-(GGSG)_5_-mCherry/GFP-His were generated by subcloning into the pHis-parallel1 vector using PCR amplification and Gibson Assembly (Sheffield et al. 1999; Gibson et al. 2009). The linker sequence (GGSG)_5_ consists of five tandem repeats of the GGSG tetrapeptide. Expression plasmids encoding His-SUMO-fused C-terminal truncation constructs encompassing residues 360-414 or 379-414 (His-SUMO-TDP-43-360-414 and His-SUMO-TDP-43-379-414, respectively) were generated in the pETDuet vector (Novagen) using PCR amplification and Gibson Assembly (Gibson et al. 2009). ALS-associated mutations and methionine-to-serine substitutions were introduced into the corresponding plasmids using QuikChange site-directed mutagenesis.

The resulting plasmids were transformed into *Escherichia coli* BL21(DE3) competent cells by heat-shock transformation. A single colony was inoculated into 10 mL of LB medium and cultured overnight at 37 °C. The overnight culture was then used to inoculate 1 L of LB medium and grown at 37 °C with shaking. When the culture reached an OD_600_ of approximately 0.8, protein expression was induced by the addition of 0.5 mM isopropyl β-D-1-thiogalactopyranoside (IPTG). The cultures were subsequently incubated at 16 °C for 20 h before harvest. Cells were collected by centrifugation at 4,000 × g for 15 min at 4 °C, washed once with phosphate-buffered saline (PBS), and stored at −80 °C until use.

His-tagged TDP-43 LCD was purified under denaturing conditions as described previously (Kato et al. 2012). All other proteins were purified under semi-denaturing conditions as described previously (Kato et al. 2017). Briefly, harvested cells were resuspended in lysis buffer containing 20 mM Tris-HCl (pH 7.5), 500 mM NaCl, 10 mM β-mercaptoethanol (BME), 0.1 mM phenylmethylsulfonyl fluoride (PMSF), and 2 M urea. Cells were lysed by the addition of lysozyme (0.2 mg/mL final concentration) followed by sonication on ice. Cell debris and insoluble material were removed by centrifugation at 48,000 × g for 1 hour at 4 °C. The resulting supernatant was loaded onto a Ni-NTA affinity column (Qiagen). After washing the column with 250 mL of wash buffer containing 20 mM Tris-HCl (pH 7.5), 500 mM NaCl, 10 mM BME, 0.1 mM PMSF, 2 M urea, and 20 mM imidazole, bound proteins were eluted with elution buffer containing 20 mM Tris-HCl (pH 7.5), 500 mM NaCl, 10 mM BME, 0.1 mM PMSF, 2 M urea, and 250 mM imidazole. The eluted proteins were concentrated using Amicon Ultra centrifugal filters (Merck Millipore) and further purified by size-exclusion chromatography on a HiLoad Superdex 200 column (Cytiva) equilibrated with S-200 buffer containing 20 mM Tris-HCl (pH 7.5), 200 mM NaCl, 1 mM dithiothreitol (DTT), and 2 M urea. Eluted fractions were analyzed by SDS-PAGE, and fractions containing intact fluorescent proteins were pooled, flash-frozen in liquid nitrogen, and stored at −80 °C until use. All purified proteins were used within one month of purification.

### Hydrogel-binding assay

The hydrogel-binding assay was performed as described previously (Kato et al. 2017). Briefly, purified TDP-43 LCD-(GGSG)_5_-mCherry-His was dialyzed against gelation buffer containing 25 mM Tris-HCl (pH 7.5), 200 mM NaCl, 1 mM DTT, and 0.5 mM EDTA. After overnight dialysis, insoluble aggregates were removed by centrifugation at 8,000 × g for 1 min, followed by brief sonication. The protein solution was then concentrated to approximately 20 mg/mL. Aliquots (0.5 μl) of the concentrated protein solution were dispensed onto glass-bottom dishes and incubated at 4 °C for 3–5 days to allow hydrogel formation.

Purified TDP-43 LCD-(GGSG)5-GFP-His wild-type and mutant proteins were diluted to 0.5 μM in gelation buffer. Insoluble material was removed by centrifugation at 8,000 × g for 10 min, and the resulting supernatant was further passed through a 0.22-μm centrifugal filter to eliminate residual aggregates. The clarified GFP-tagged protein samples were then added to the preformed hydrogels and incubated overnight at 4 °C. Fluorescence images were acquired using a ZEISS LSM 980 Airyscan confocal microscope. GFP fluorescence intensity within the hydrogels was quantified using ImageJ software as described previously (Kato et al. 2017). Measurements from three independent experiments were averaged, and the results are presented as mean ± standard deviation.

### Thioflavin T assay for TDP-43 LCD

Wild-type His-TDP-43 LCD and its mutant derivatives were diluted to 300 μM in a buffer containing 25 mM Tris-HCl (pH 7.5), 200 mM NaCl, and 6 M guanidine hydrochloride (GdnHCl) to standardize the protein concentration. For the analysis of ALS-associated mutants, the samples were then diluted to a final protein concentration of 4 μM in thioflavin T (ThT) assay buffer containing 25 mM Tris-HCl (pH 7.5), 200 mM NaCl, 1 mM BME, and 40 μM ThT. For the analysis of M-to-S mutants, the samples were diluted to a final protein concentration of 10 μM in the ThT assay buffer containing 30 μM ThT. Immediately after dilution and vortex mixing, the samples were transferred to a 96-well black fluorescence microplate at a volume of 100 μL per well. The plate was sealed with a clear film to prevent evaporation during incubation. ThT fluorescence was monitored for 20–48 h using a SpectraMax iD3 (Molecular Devices) or an Infinite F Nano+ (TECAN) plate reader. Measurements were recorded at 15-min intervals with 20 s of shaking before each reading. Wells containing ThT assay buffer without protein were used as blanks for background subtraction. ThT fluorescence was measured in three to six replicate wells. For each well, baseline fluorescence was calculated as the mean ThT signal during the first hour of the assay. Lag time was defined as the elapsed time from the start of the measurement until the ThT fluorescence signal first reached 200% of the mean baseline fluorescence value. Lag times calculated for individual wells were averaged and are presented as mean ± standard deviation.

### *In vitro* fibrillization of CCR-containing C-terminal constructs

Purified His-SUMO-TDP-43-360-414 and His-SUMO-TDP-43-379-414 were diluted to 1 mg/mL in gelation buffer, and Ulp1 SUMO protease was added at an enzyme-to-substrate ratio of 1:250. The mixtures were incubated with gentle rotation at 4 °C overnight. Following cleavage, the samples were concentrated to approximately 2 mL and subjected to size-exclusion chromatography to separate the cleaved TDP-43 fragments from the His-SUMO tag and Ulp1. The gel filtration column was equilibrated with gel filtration buffer containing 20 mM Tris-HCl (pH 7.5), 200 mM NaCl, 1 mM DTT, and 2 M GdnHCl. Fractions containing the cleaved TDP-43 C-terminal fragments were pooled and concentrated to approximately 30 mg/mL. The purified fragments were then diluted to 3 mg/mL in the gel filtration buffer with 6 M GdnHCl and dialyzed overnight against fibrillization buffer consisting of 20 mM Tris-HCl (pH 7.5), 200 mM NaCl, 10 mM BME, 0.5 mM EDTA, and 0.1 mM PMSF. After dialysis, the protein samples were transferred to microcentrifuge tubes and incubated at 4 °C for 1 week to allow fibril formation. Brief sonication was performed every 3 days to facilitate fibrillization. Fibril formation was examined by negative-stain transmission electron microscopy as described previously (Kato et al. 2021).

### ThT analysis of CCR-containing fragments

Purified TDP-43 360-414 and 379-414 fragments were diluted to a final concentration of 300 μM in the ThT assay buffer containing 30 μM ThT. ThT fluorescence was monitored using a Cytation 5 plate reader (Agilent) with measurements recorded at 15-min intervals and 5 min of shaking before each reading. Each trace represents the mean fluorescence intensity ± standard deviation from six replicate wells.

### Statistical analyses of the hydrogel-binding assays and ThT assays

Statistical comparisons of multiple mutants were performed by one-way analysis of variance (ANOVA) followed by Dunnett’s multiple-comparisons test, using wild-type TDP-43 LCD as the reference group for the data shown in Figures 2, 4, and 5. These analyses were performed using the online Stats Calculator program (https://www.statscalculators.com/calculators/hypothesis-testing/dunnetts-test-calculator). For the ThT assay data shown in Figure 3, lag times were analyzed using Kaplan-Meier survival analysis to account for right-censored data (>24 h). Pairwise comparisons between each mutant and wild-type TDP-43 were performed using the log-rank test. Multiple comparisons were adjusted using the Bonferroni correction. These analyses were conducted using the OASIS 2 platform (https://sbi.postech.ac.kr/oasis2/). Differences were considered statistically significant at P < 0.05.

## Acknowledgments

We thank Motomasa Tanaka of RIKEN and Naruhiko Sahara of Niigata University for valuable discussions regarding this research. We thank Steven McKnight of the University of Texas Southwestern Medical Center for providing the expression plasmids for the original TDP-43 LCD. We also thank Jinge Gu for providing purified Upl1 protein. This research was supported by AMED-CREST Grant Number 26gm1410009h0005, the MEXT Quantum Leap Flagship Program (Q-LEAP) Grant Number JPMXS0120330644, and JSPS KAKENHI Grant Number 23K23857. M.K. was additionally supported by NIGMS Grant 5R35GM130358-08.

## Author contributions

N.M, S.-Y.K. and M.K. designed research; N.M, S.-Y.K. Y.K. and M.K. performed research; N.M and S.-Y.K. analyzed data; and N.M, S.-Y.K. and M.K. wrote the paper.

The authors declare no competing interests.

The data supporting the findings of this study are available from the corresponding author upon reasonable request.

## Supplementary Figures

**Supplementary Figure 1.**
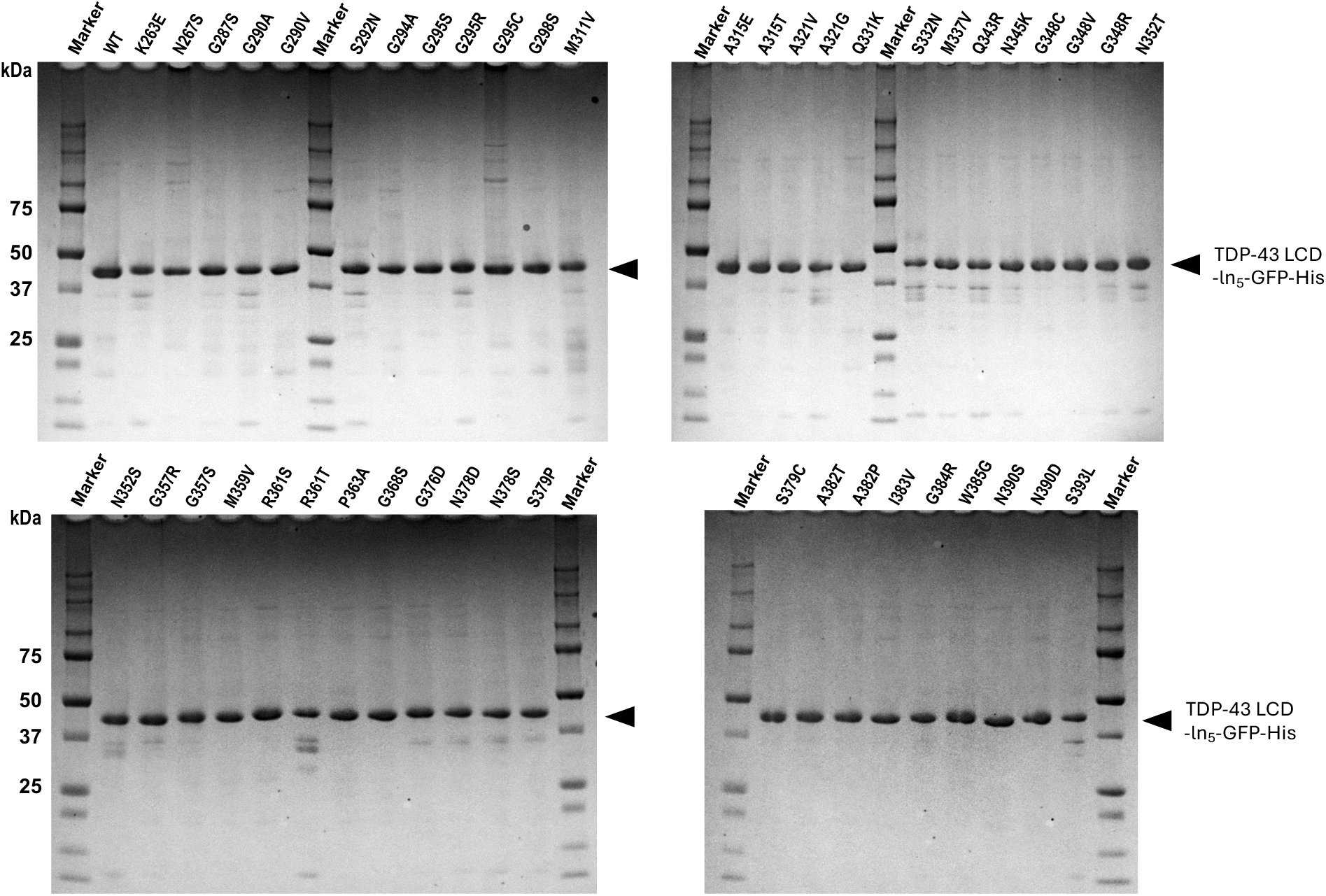
Expression and purification of TDP-43 LCD ALS mutants. Purified GFP-tagged wild-type and ALS-associated mutant TDP-43 LCD proteins were analyzed by SDS-PAGE followed by Coomassie Brilliant Blue staining. Proteins were purified by Ni-affinity and gel-filtration chromatography.

**Supplementary Figure 2.**
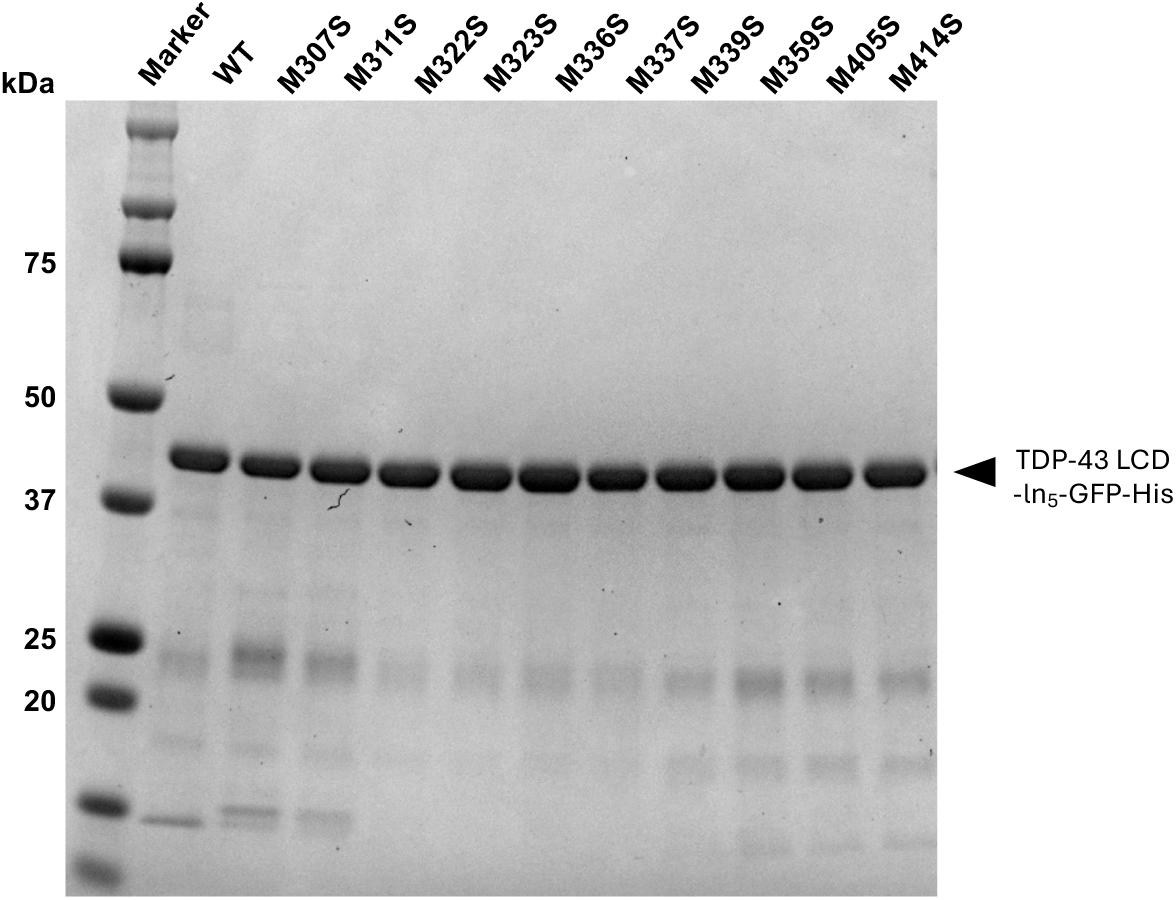
Purification of methionine-to-serine mutants. SDS-PAGE analysis of purified wild-type and methionine-to-serine mutant TDP-43 LCD-EGFP proteins used in this study.

**Supplementary Figure 3.**
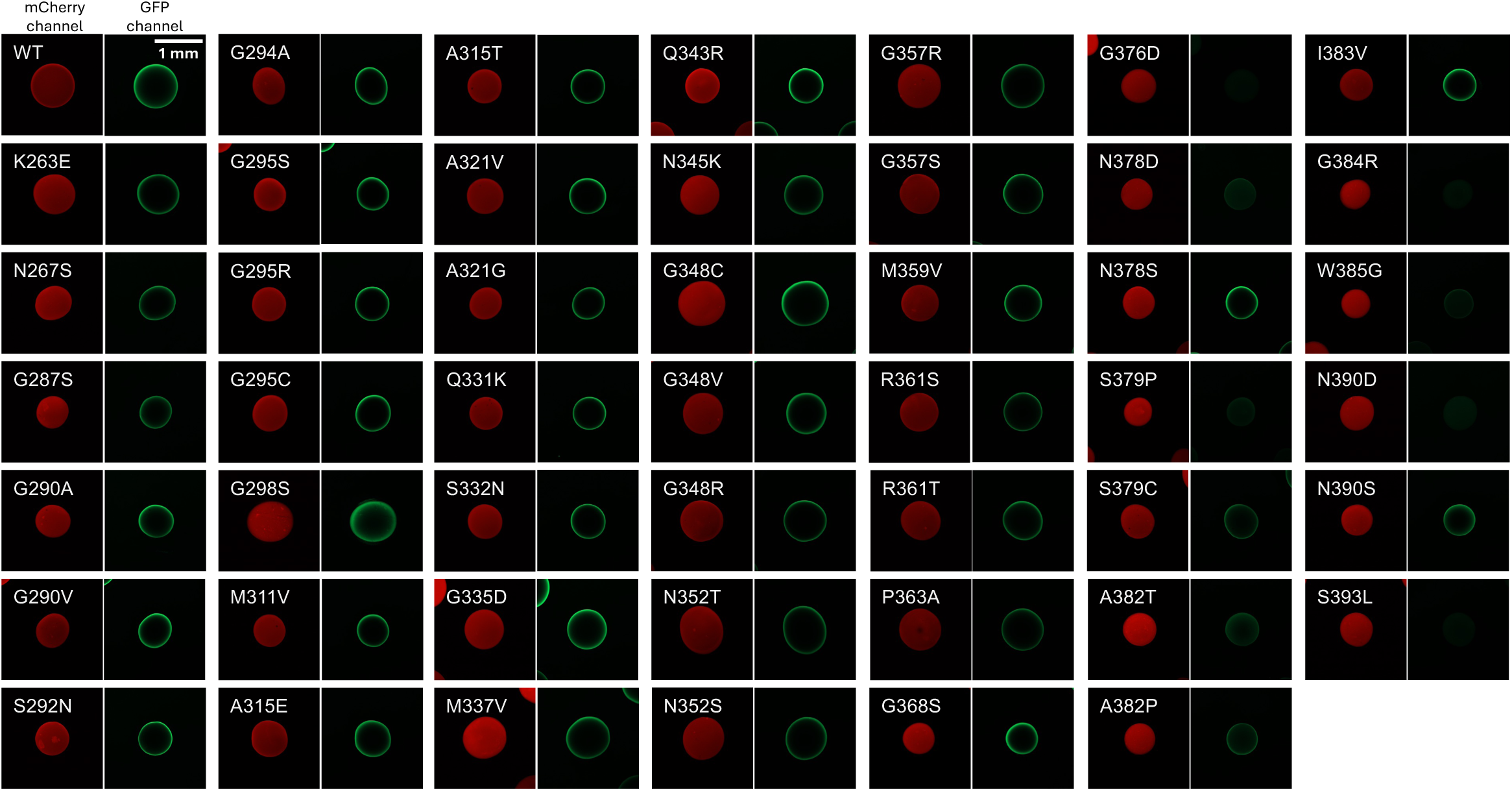
Representative images of the hydrogel-binding assays for ALS-associated mutants of TDP-43 LCD. Representative confocal fluorescence images showing incorporation of individual ALS-associated GFP-TDP-43 LCD mutants into preformed mCherry-TDP-43 LCD hydrogels. Reduced incorporation was particularly evident for several mutations located within the CCR. All GFP images were displayed using identical intensity settings. Scale bar, 1 mm.

**Supplementary Figure 4.**
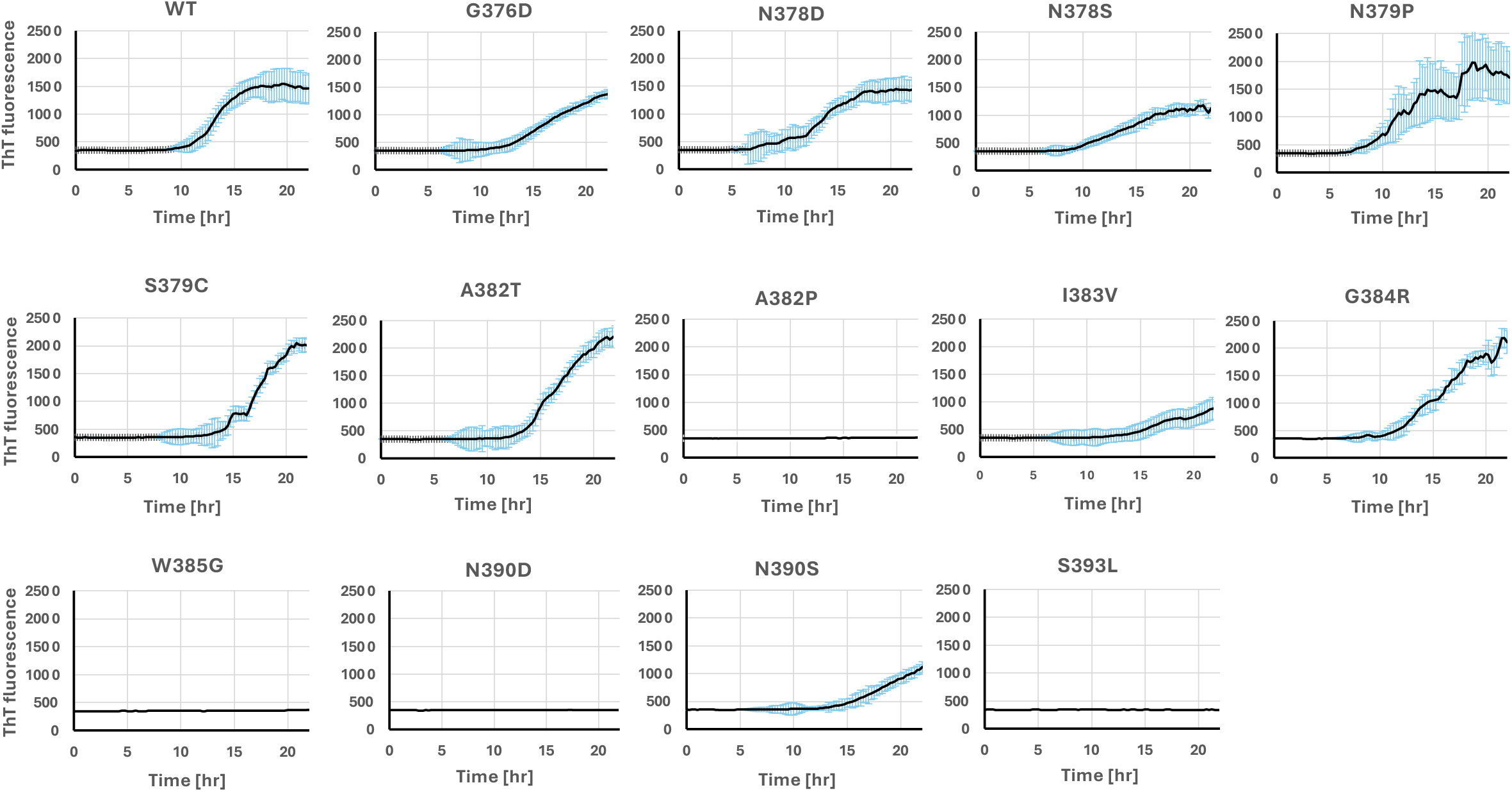
Thioflavin T fluorescence kinetics of ALS-associated CCR mutants. ThT fluorescence was monitored at 15-min intervals under the indicated assay conditions. Each curve represents the mean fluorescence intensity obtained from six replicate wells. The cyan bars indicate standard deviation. Lag times shown in Figure 3 were determined as described in Materials and Methods.

**Supplementary Figure 5.**
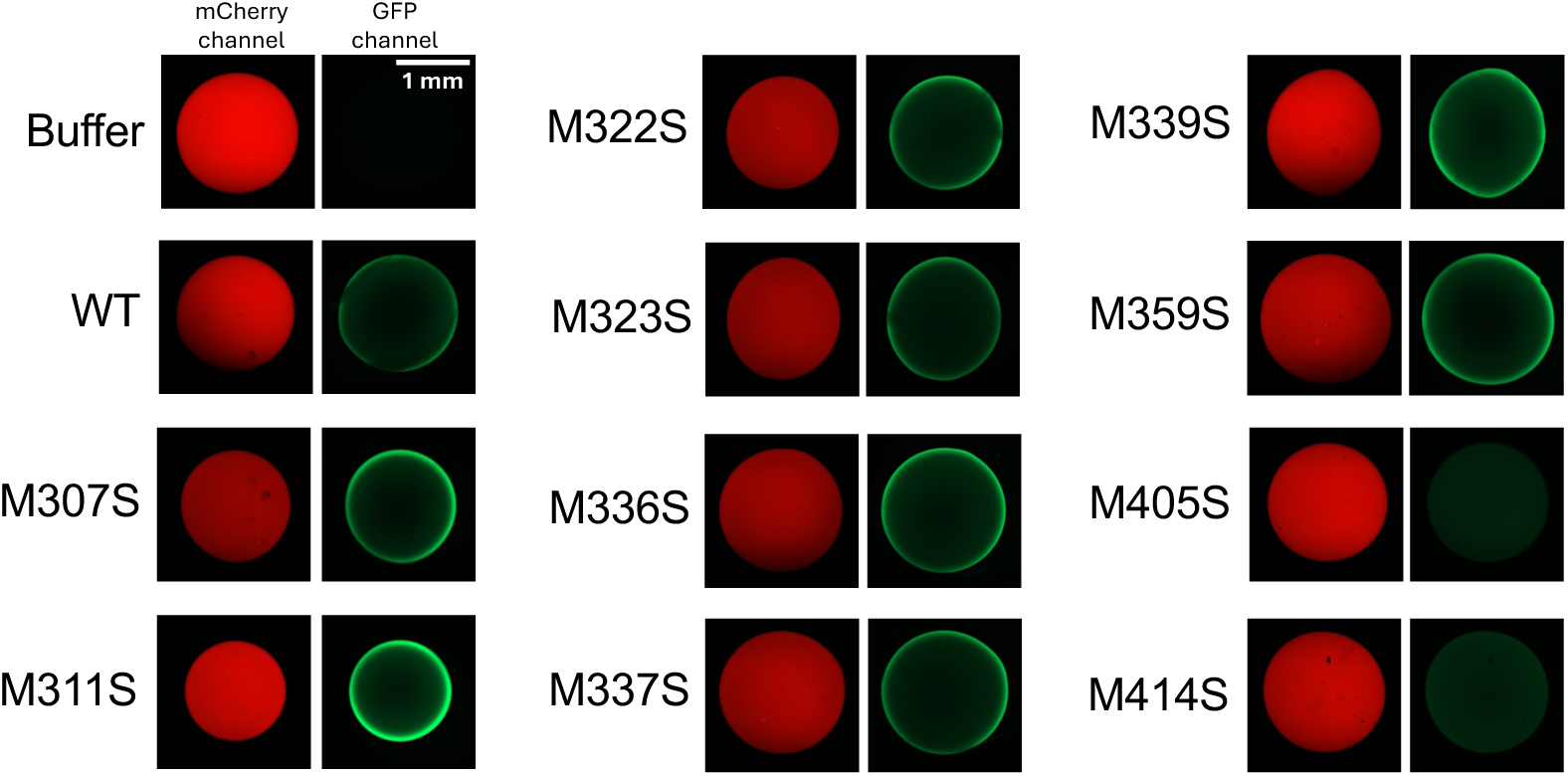
Representative hydrogel-binding assays of methionine-to-serine mutants. Representative confocal fluorescence images showing incorporation of methionine-to-serine mutant GFP-TDP-43 LCD proteins into preformed mCherry-TDP-43 LCD hydrogels. Reduced hydrogel incorporation was observed for M405S and M414S. Scale bar, 1 mm.

**Supplementary Figure 6.**
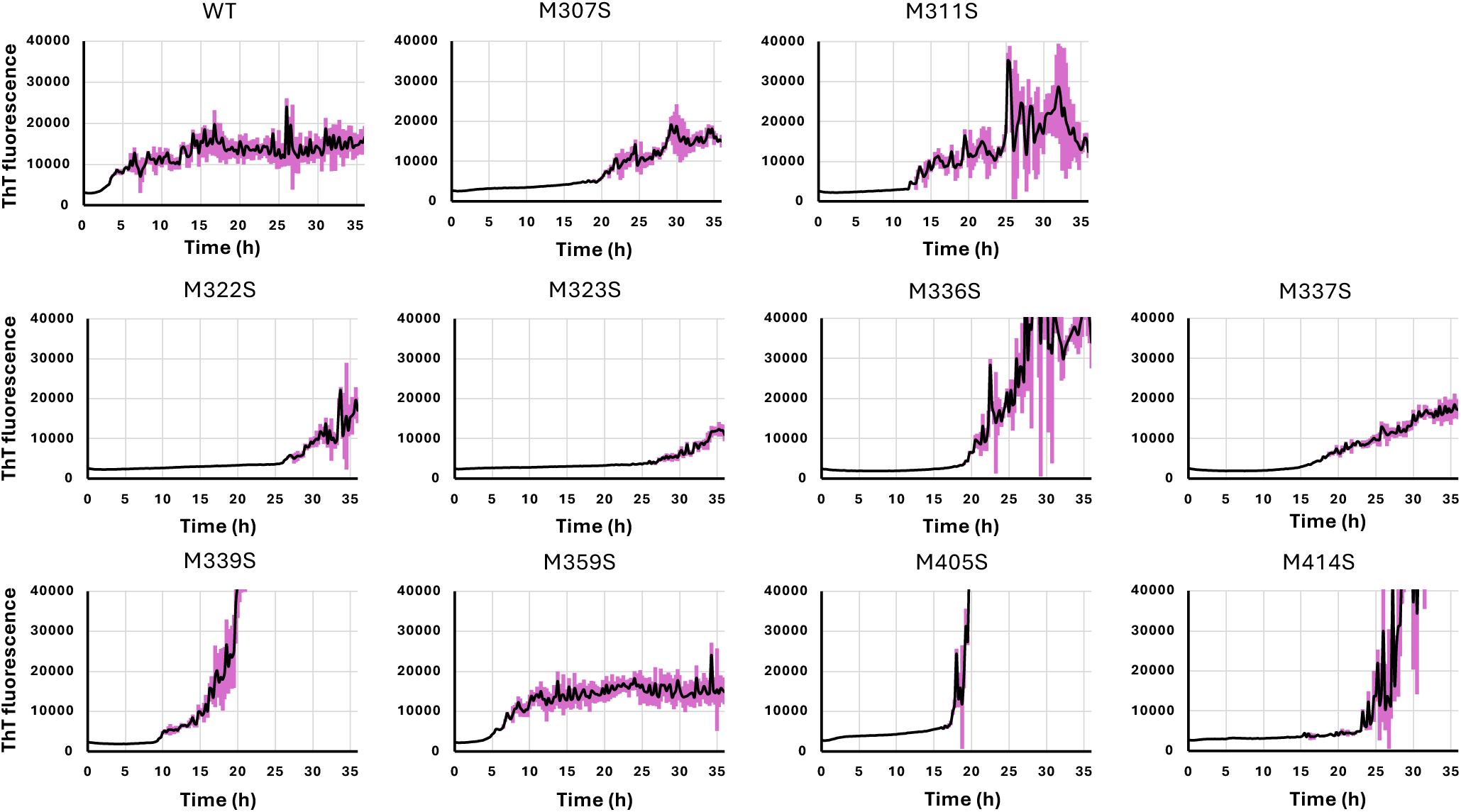
ThT fluorescence kinetics of wild-type and methionine-to-serine mutant TDP-43 LCD proteins. ThT fluorescence was monitored at 15-min intervals under the assay conditions indicated in Materials and Methods. Each curve represents the mean fluorescence intensity obtained six replicate wells. The purple bars indicate standard deviation. Lag times shown in Figure 5 were determined as described in Materials and Methods.

